# A hybrid model of the within-host dynamics post-infection with Legionnaires’ disease

**DOI:** 10.1101/2025.08.31.673357

**Authors:** Nyall Jamieson, Christiana Charalambous, David M. Schultz, Ian Hall

**Affiliations:** Department of Mathematics, The University of Manchester, Upper Brook Street, Manchester, M13 9PL, United Kingdom; Centre for Atmospheric Sciences, Department of Earth and Environmental Sciences, and Centre for Crisis Studies and Mitigation, The University of Manchester, Oxford Road, Manchester, M13 9PL, United Kingdom

**Keywords:** branching process, first passage time distribution, incubation period, Legionnaires’ disease, stochastic differential equation, uncertainty quantification

## Abstract

Understanding the incubation period of Legionnaires’ disease is vital for accurate source-term identification. Traditionally, researchers estimate the dose-dependent incubation period from human outbreak data, but this method suffers from the inability to estimate the exposure dose retrospectively for each case. This challenge limit the precision of incubation-period analysis using human case data. Existing within-host models, such as ordinary differential equation (ODE)-based and discrete-event stochastic approaches, estimate the dose-dependent incubation period of Legionnaires’ disease. However, discrete-event models, while useful, are so computationally costly that the within-host dynamics must be simplified to solely the *Legionella* and macrophage interactions. This simplification makes the computation feasible, but precludes cytokine interactions and adaptive immune response modelling. In this paper, we develop a new approach to model the within-host dynamics of Legionnaires’ disease that focuses on reducing computational cost while maintaining accuracy. Specifically, we propose a hybrid framework that integrates and improves upon existing ODE and discrete event within-host models of Legionnaires’ disease. By integrating the previously developed ODE and discrete-event stochastic models with stochastic differential equation (SDE) models, we create a unified system that adapts dynamically throughout the infection process. We quantify the points at which each model becomes the optimal tool for describing the infection, resulting in a flexible simulation of disease dynamics. Our hybrid model aligns with observed human incubation-period data and is the first framework of its kind in this context. This advancement offers a more robust platform for testing additional biological assumptions and improving our understanding of Legionnaires’ disease.

## 1. Introduction

*Legionella* is a gram-negative bacterium that causes Legionnaires’ disease, a serious condition due to the bacteria’s ability to evade the human immune response and infect phagocytic cells in the lungs. This bacteria transmits through inhalation of aerosolized water droplets contaminated with *Legionella*, which often originate from water sources such as cooling towers, humidifiers, and whirlpool spas. Once inhaled, *Legionella* enter the lungs and initiate infection as phagocytic cells attempt to engulf the bacteria. Inside these cells, *Legionella* proliferates until the phagocyte ruptures, releasing the bacteria back into the lungs and enabling further infection. This cycle of infection and rupture of cells can lead to severe lung damage and systemic illness. Symptoms of this disease include fever, headache, lethargy, hemoptysis, respiratory failure, multiple organ failure, and fatal pneumonia. According to [1], Legionnaires’ disease has a mortality rate of 5–10% in the general population and 40–80% in untreated immunosuppressed individuals. Typically, the disease predominantly appears in older males, as 75–80% of cases are over 50 years and 60–70% of cases are male.

As a result, understanding Legionnaires’ disease is crucial for public health, environmental monitoring, and risk management. One key aspect of this understanding is the incubation period of Legionnaires’ disease, which is the delay in time between exposure to *Legionella* and onset of symptoms. A reliable understanding of this delay is crucial for effective disease control; by correctly quantifying the incubation period, public health authorities can reliably estimate the exposure window, identify possible sources of contamination, and implement targeted interventions to prevent further cases [15]. Moreover, understanding the incubation period enables the timely identification of the at-risk population, ensuring faster diagnosis and treatment, which reduces the severity of symptoms and saves lives.

Mathematical models are useful tools for describing various stages of *Legionella*’s life cycle and transmission pathway. For instance, different models have been used to describe population growth in water and biofilms [18, 20, 23], aerosolization [14, 22, 25], and within-host dynamics [16]. The importance of choosing the appropriate modelling approach has been highlighted [16], which suggests that, in the within-host case, deterministic models may be less effective than stochastic models in certain contexts, and vice versa. Initially, the stochastic nature of within-host *Legionella* population growth poses challenges for ODEs. These models become effective only after the population surpasses a certain threshold, as stochastic fluctuations dominate at lower population levels. Therefore, when employed in the early stages of infection, an ODE model fails to account for the stochastic timing of phagocytic and rupture events, as well as the variable number of *Legionella* released in a rupture event. Additionally, an ODE model describing exponential bacterial growth, as in [16], does not allow for an individual to recover from infection. Moreover, employing an ODE does not accurately estimate the uncertainty associated with the incubation period and fails to capture the variability observed in human outbreaks [4, 6, 8, 10, 11, 13]. Therefore, although ODEs are computationally cheap to run, they introduce several issues that hinder the accurate representation of important features of the biological process. However, as the *Legionella* population in the lungs grows, the frequency of rupture events increases over time. Eventually, due to the law of large numbers, the total number of *Legionella* released from these events approaches the expected rupture size. The averaging effect ensures that ODEs may accurately describe the within-host dynamics when the bacterial population is sufficiently large [5, 16].

To overcome the limitations of ODEs in modelling the early-stage *Legionella* infection dynamics, a discrete-event stochastic model has been employed to accurately represent the random timing of phagocytic and rupture events, as well as the variable number of *Legionella* released during macrophage rupture [16]. The stochastic model accounts for both the variability in the rupture sizes and the inherent randomness in event timing. As a result, this model yields incubation periods that align closely with human outbreak data [4, 6, 8, 10, 11, 13]. Unfortunately, as the *Legionella* population increases, the computational cost of stochastic simulations becomes prohibitive. This makes them impractical for simulating the entire infection process or conducting large-scale sensitivity analyses [5]. Furthermore, stochastic simulations, by their nature, often do not provide explicit mathematical insight into the true underlying distribution of the *Legionella* population at any given time [5]. To balance accuracy and computational feasibility, a natural solution may be to adopt a hybrid approach that dynamically transitions between modeling frameworks based on the system’s state. Initially, a discrete-event stochastic model may be used when populations are small and stochasticity dominates. Once the populations grow large enough for fluctuations to average out and deterministic dynamics emerge, the hybrid model may switch to an ODE. This strategy ensures that the model remains computationally tractable while retaining sufficient detail to capture key system behaviours at each stage. Such hybrid models have been successfully applied in epidemic modeling in the context of SIR models [17]. These hybrid models switch between deterministic and stochastic models to describe disease transmission through a population by counting the number of infected individuals [3, 5, 9, 26]. These models provide a scalable means of describing disease dynamics across different population sizes and phases of an outbreak.

To extend this hybrid modelling approach, stochastic differential equations (SDEs) with Gaussian noise emerge as a natural bridge between discrete-event stochastic models and ODE models. While ODEs provide a deterministic framework and discrete-event stochastic models offer realism with their stochasticity, SDE models are computationally fast, and allow bacterial populations to grow approximately according to the mean growth curve with a noise term for stochasticity [2]. Additionally, due to the stochastic noise term, SDE models allow for bacterial populations to reach zero where extinction occurs [2]. These two properties suggest that SDE models may possess the strengths of both the discrete-event stochastic and ODE models. However, when the extracellular *Legionella* population is low relative to the expected rupture size, Gaussian SDE models may fail to capture the large jumps in the extracellular *Legionella* population (a visualisation of this issue is provided in Fig. 2a). This indicates that like an ODE model, a Gaussian SDE model cannot be employed immediately after infection.

Therefore, employing a Gaussian SDE model as an intermediary model between the discrete-event stochastic and ODE models may provide a smoother, computationally cheaper and accurate alternative to the discrete-event-to-ODE hybrid model framework.

In this paper, we propose a novel hybrid framework to describe *Legionella* population dynamics, incorporating three distinct modelling approaches to capture different phases of the infection process. Our approach first combines a multi-type branching process (MTBP) model for the early post infection phase with an ODE model for the later phase. Additionally, we introduce a Gaussian SDE model as an intermediary between the MTBP and ODE models. This intermediary stage allows for a smoother transition by capturing intermediate fluctuations that the ODE alone cannot. We identify the conditions under which each model provides an appropriate approximation of the within-host dynamics as we quantify the transition times accordingly. To the best of our knowledge, this is the first hybrid model for bacterial infection dynamics that explicitly integrates a Gaussian SDE as an intermediary step. Our work provides a novel framework for systematically determining when different modelling approaches become appropriate. By developing and implementing this approach, we aim to offer an improved framework for analysing the incubation period of Legionnaires’ disease and demonstrate the broader applicability of multi-stage hybrid models in infectious disease modelling.

## 2. Methods

In this section, we discuss various methods for modelling the within-host dynamics following infection with Legionnaires’ disease. We first present the continuous-time Markov chain (CTMC) model developed in [16] in Section 2.1 and then derive a MTBP approximation of this model in Section 2.2. Next, we propose a jump SDE in Section 2.3 that captures the large variable increase in the *Legionella* population following rupture events. This model also describes the early within-host dynamics that occur immediately after infection.

Following this, in Section 2.4 we derive an SDE model of the within-host dynamics, which consists of a deterministic drift component added to a Wiener process (Gaussian noise). We refer to this as the Gaussian SDE model. Finally, in Section 2.5 we describe the ODE model for the within-host dynamics of Legionnaires’ disease developed in [16]. We provide a discussion on the shortcomings of Gaussian SDE and ODE models when applied from the beginning of infection in Section 2.6. Following this discussion, in Section 2.7 we define two first passage times: the first marks when a Gaussian SDE model accurately approximates the MTBP dynamics 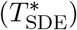, and the second marks when an ODE model does the same 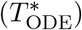. These first passage times are essential for developing a hybrid framework of the within-host dynamics of Legionnaires’ disease.

We consider two approaches for developing a hybrid model of the infection process. First, we derive a hybrid model with one transition at the first passage time 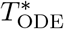. At this time, a MTBP model switches to an ODE model once an ODE model provides an accurate approximation of the MTBP dynamics. Second, we derive a hybrid model as above, with a Gaussian SDE as an intermediary between the MTBP and ODE models. In this scenario, the MTBP model switches to a Gaussian SDE model at the first passage time 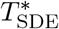, and then to an ODE at 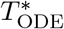(Fig. 1). For each hybrid model, illness occurs at time 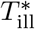, once the extracellular *Legionella* population reaches a threshold *L*_*T*_ = 50661 *Legionella* within the lungs [16].

**Figure 1.**
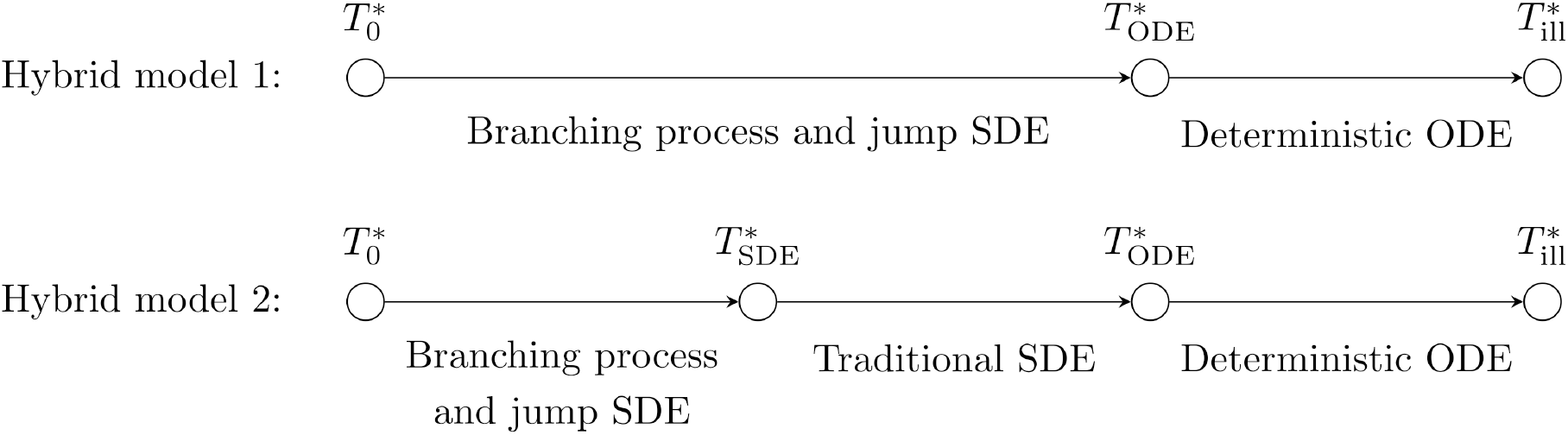
A visual representation of the two hybrid models. Each model starts with infection at the 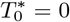. Between 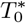 and 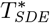, we expect only the MTBP model and jump SDE model to accurately describe the within-host dynamics. Between 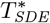 and 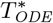, we expect the Gaussian SDE model to also provide an accurate simplification of the MTBP model. Additionally, from 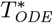 to 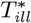 we expect the ODE model to also provide an accurate simplification of the MTBP model. 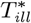 represents the time at which a extracellular Legionella population reaches the threshold number residing within the lungs. Here we assume an onset of symptoms occurs within the individual at 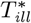.

### 2.1. Within-host CTMC model

We consider a stochastic model for the within-host dynamics of an individual infected with Legionnaires’ disease [16]. Once an individual becomes infected, the bacteria reside within the lungs, where one of three key events may occur. Macrophages within the lungs may attempt to engulf the *Legionella*, leading to two possible outcomes. In the first case, macrophages kill the *Legionella* during this process, causing individual *Legionella* to die at rate *β*. Alternatively, the *Legionella* may survive phagocytosis at rate *α*, then reside within the macrophage, and reproduce. Finally, infected macrophages rupture, releasing their intracellular *Legionella* population back into the lungs. In this event, individual infected macrophages rupture at rate *λ*. During a rupture event, the number of intracellular *Legionella* released follows a negative binomial (NB) distribution with mean *G* and overdispersion parameter *r*. Moreover, to fit within a Markovian framework, we treat the time until macrophage rupture and the resulting intracellular *Legionella* populations as independent from one another [16]. The three types of interactions between *Legionella* and macrophages may persist until the extracellular *Legionella* population reaches a threshold population *L*_*T*_, at which point symptoms appear.

We consider a two-dimensional stochastic process ***Z***(*t*) = (*L*(*t*), *M* (*t*)), in which we record the number of extracellular *Legionella L*(*t*) and infected macrophages *M* (*t*) at a time *t*. For the three key events described above, the transitions between states occur at the following rates:

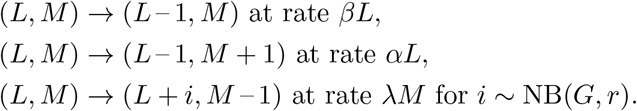

### 2.2. Multi-type branching process approximation

The CTMC model may be described in a multi-type branching process (MTBP) framework [7]. We consider the branching process with two types: extracellular *Legionella* and infected macrophages. After each of the three key events within the lungs, an individual dies, and a number of individuals of either type are born.

Individual *Legionella* die at rate *ω*_*L*_ = *α* + *β*, whereas individual infected macrophages die at rate *ω*_*M*_ = *λ*. In this framework, two possibilities may occur when an extracellular *Legionella* dies. First, no individual is born, with probability *β/*(*α* + *β*). Second, an infected macrophage is born, with probability *α/*(*α* + *β*). Similarly, when an infected macrophage dies, a population of extracellular *Legionella* is born. The size of this population follows a negative binomial distribution with mean *G* and overdispersion parameter *r*. We derive the progeny generating function *P*_*i*_ for the number of offspring produced when an individual of type *i* ∈ {*L, M*} dies as follows:

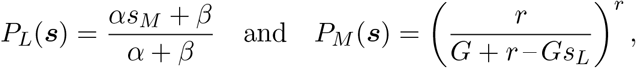

where ***s*** = (*s*_*L*_, *s*_*M*_ ) ∈ (0, 1)^2^. For brevity, we define ***P*** (***s***) = (*P*_*L*_(***s***), *P*_*M*_ (***s***)). Next, we consider the probability generating function for ***Z***(*t*), conditional on the infection process beginningwith a single individual of type *i* ∈ {*L, M*}:

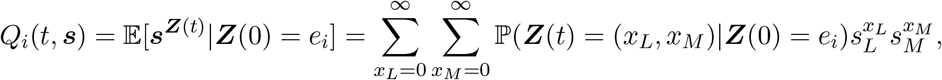

where *e*_*i*_ represents the vector with one in the *i*^th^ component and zero everywhere else. Therefore, *e*_*i*_ represents an initial population of one individual of type *i* ∈ {*L, M*}. Further, we define ***Q***(*t*, ***s***) = (*Q*_*L*_(*t*, ***s***), *Q*_*M*_ (*t*, ***s***)) as the vector form of the probability generating function. In the MTBP context, these probability generating functions satisfy ∂*Q*_*i*_*/*∂*t* = – *ω*_*i*_*Q*_*i*_(*t*, ***s***) + *ω*_*i*_*P*_*i*_(*Q*(*t*, ***s***)) for *i* ∈ {*L, M*} [7]. Therefore, the within-host dynamics satisfy the following system of partial differential equations:

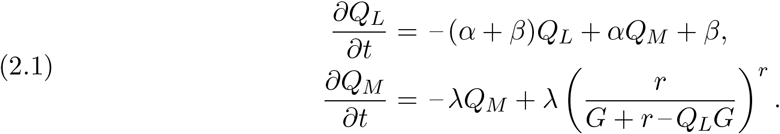

The solution *q*_BP_(*t*) = *Q*_*L*_(*t*, **0**) of (2.1) gives the probability that a branching process initiated by a single extracelluar *Legionella* goes extinct by time *t*. From this solution, we define the instantaneous probability that extinction occurs as *p*(*t*) = *dq*(*t*)*/dt*. Next, we define a matrix **Ω**, in which the entry in the *i*^th^ row and *j*^th^ column equals 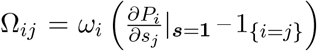. An element in the *i*^th^ row and *j*^th^ column of this matrix represents the expected rate of change in the type *j* ∈ {*L, M*} population due to the death of a type *i* ∈ {*L, M*} individual. In our scenario, we define **Ω** as follows:

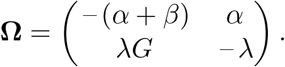

We define the mean vector at time *t* as ***m***(*t*) = (𝔼 [*L*(*t*)], 𝔼 [*M* (*t*)]), which satisfies the following equation:

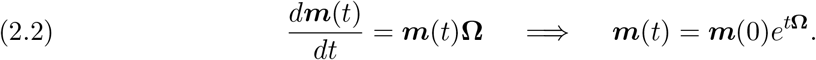

Thus the solution of (2.2), with the derivation left to the Appendix, is as follows:

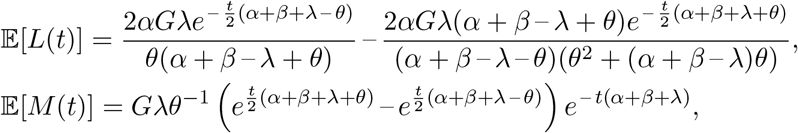

where 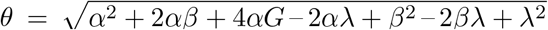. Further, the variance matrix for the MTBP, starting with a single individual of type *i* ∈ {*L, M*}, satisfies 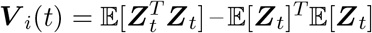. Additionally, the variance matrix also satisfies the following differential equation [7]:

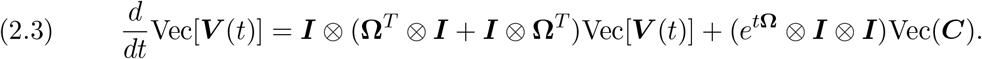

In (2.3), ***I*** is the identity matrix, ⊗ is the Kronecker product of matrices, the Vec operator stacks columns of a matrix to form a single column vector, and Vec(***C***) is the stacked column vector of Vec(*d****V*** _*j*_(0)*/dt*) for *j* =∈ {*L, M*}. In addition, we provide calculations involving the variance in the Appendix.

### 2.3. Jump SDE

We derive a jump process SDE to approximate the MTBP. Specifically, we approximate the MTBP with a discrete-time stochastic process that counts the number of occurrences of each event during discrete time intervals. Because the time between events in the MTBP framework is exponentially distributed, we model the number of events within a time interval using a Poisson distribution.

We consider three Poisson processes *Z*_*k*_ for *k* ∈ {1, 2, 3} to model the three key events that occur within the lungs. *Z*_1_ corresponds to the number of events in which an extracellular *Legionella* dies during phagocytosis, *Z*_2_ corresponds to the number of events in which an extracellular *Legionella* survives phagocytosis and infects a macrophage, and *Z*_3_ corresponds to the number of events in which an infected macrophage ruptures and releases a population of *Legionella* back into the lungs. Additionally, the process *Z*_4_ represents the total number of *Legionella* released back into the lungs following the *Z*_3_ macrophage rupture events. The number of *Legionella* released during each individual macrophage rupture event is independent of *Z*_3_. However, *Z*_4_ is the sum of |*Z*_3_| independent and identically distributed negative binomial processes. Each of these negative binomial processes represents the number of intracellular *Legionella* released back into the lungs during a rupture event. Because we explicitly capture the increase in the extracellular *Legionella* population following a macrophage rupture event, we assume that this model is valid immediately after infection. The jump SDE is defined as follows:

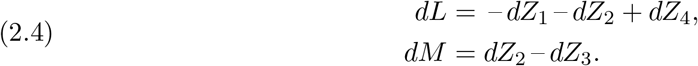

### 2.4. Gaussian SDE

We propose a Gaussian SDE to describe the within-host dynamics of Legionnaires’ disease in the intermediate stage of the process. This SDE consists of a deterministic drift term and a Gaussian diffusion term. The Gaussian noise term is used to represent the stochasticity of this within-host process and is justified as follows. In Section 2.1, we establish that the time until a rupture event follows an exponential distribution, while the rupture size follows a negative binomial distribution. Consequently, over a given time interval of length *t*, the increase in extracellular *Legionella* follows the Poisson–negative binomial compound distribution, denoted as *Z*_4_. This compound distribution has finite variance under the assumption that *λ, G* and *r* are finite.

The Gaussian noise term is used to represent the stochasticity of this within-host process. However, in small populations, the noise term does not explicitly account for the large jumps that occur when an infected macrophage ruptures. As a result, we expect that this model does not reliably describe the within-host dynamics immediately after infection, and a period of time is required before this model may approximate the MTBP dynamics. We obtain this SDE using an approach from [2], which a general method for deriving SDEs is developed. Following [2], we calculate the variance–covariance matrix ***U*** (*t*) of the within-host dynamics, which we define as follows:

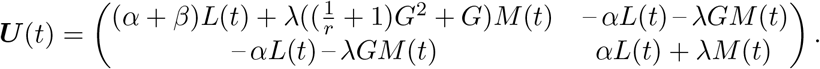

The derivation of ***U*** (*t*) is provided in the Appendix. Next, we define the square root of ***U*** (*t*) as follows:

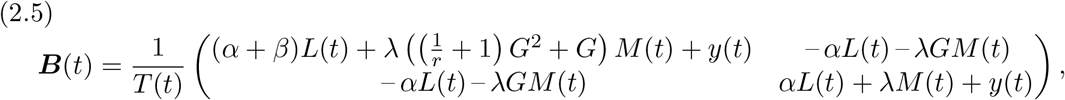

where

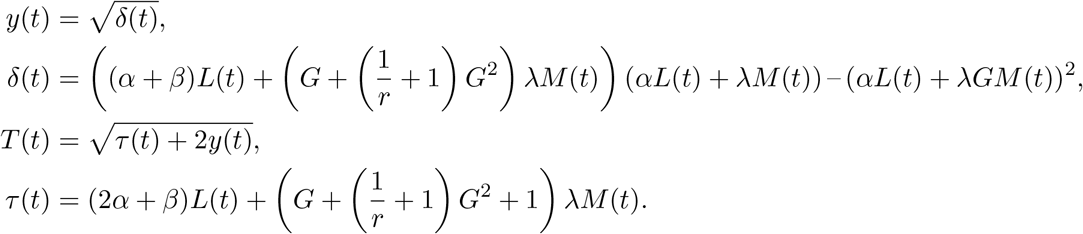

For notational simplicity, we define the column vector ***X***(*t*) = ***Z***(*t*)^*T*^. The Gaussian SDE representing the within-host dynamics is defined as follows with *d****W*** _*t*_ ∼ N(0, *t*):

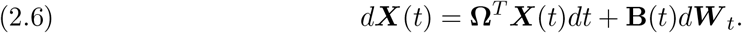

### 2.5. Deterministic system

After a sufficient period of time since infection has elapsed, the number of extracellular *Legionella* and infected macrophages within the lungs increases to a large amount. As the population grows, phagocytosis and rupture events become more frequent. This increased frequency leads to fluctuations in event timing becoming proportionally smaller around the average behavior. Under these conditions, the within-host dynamics described by the MTBP model are well approximated by the ODE model described in [16]. The system of equations is defined as follows:

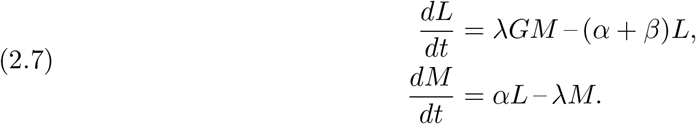

### 2.6. Justification for the hybrid framework

In the introduction, we postulated that although the SDE and ODE models may describe the within-host dynamics of Legionnaires’ disease, they cannot reliably capture these dynamics immediately after infection. In the case of ODEs, the inability to capture early infection dynamics may be more evidence due to their deterministic nature, which does not account for stochastic fluctuations present in the initial stages of infection. However, understanding how a Gaussian SDE fails to capture the variability of large jump processes within an MTBP framework is less intuitive. Although SDEs incorporate stochasticity, making them more similar to discrete-event stochastic models than ODEs are, they still fall short of capturing these large jumps. Shortly after infection, the *Legionella* population released back into the lungs following rupture events is large relative to the extracellular *Legionella* population. Incidentally, the variance term in the Gaussian SDE model cannot capture the jumps that occur when the rupture size is large relative to the extracellular *Legionella* population. A demonstration of this issue is presented in Fig. 2a, as realisations of both the SDE and discrete-event models validate the inability of the Gaussian SDE model to accurately represent large jumps in the MTBP model.

**Figure 2.**
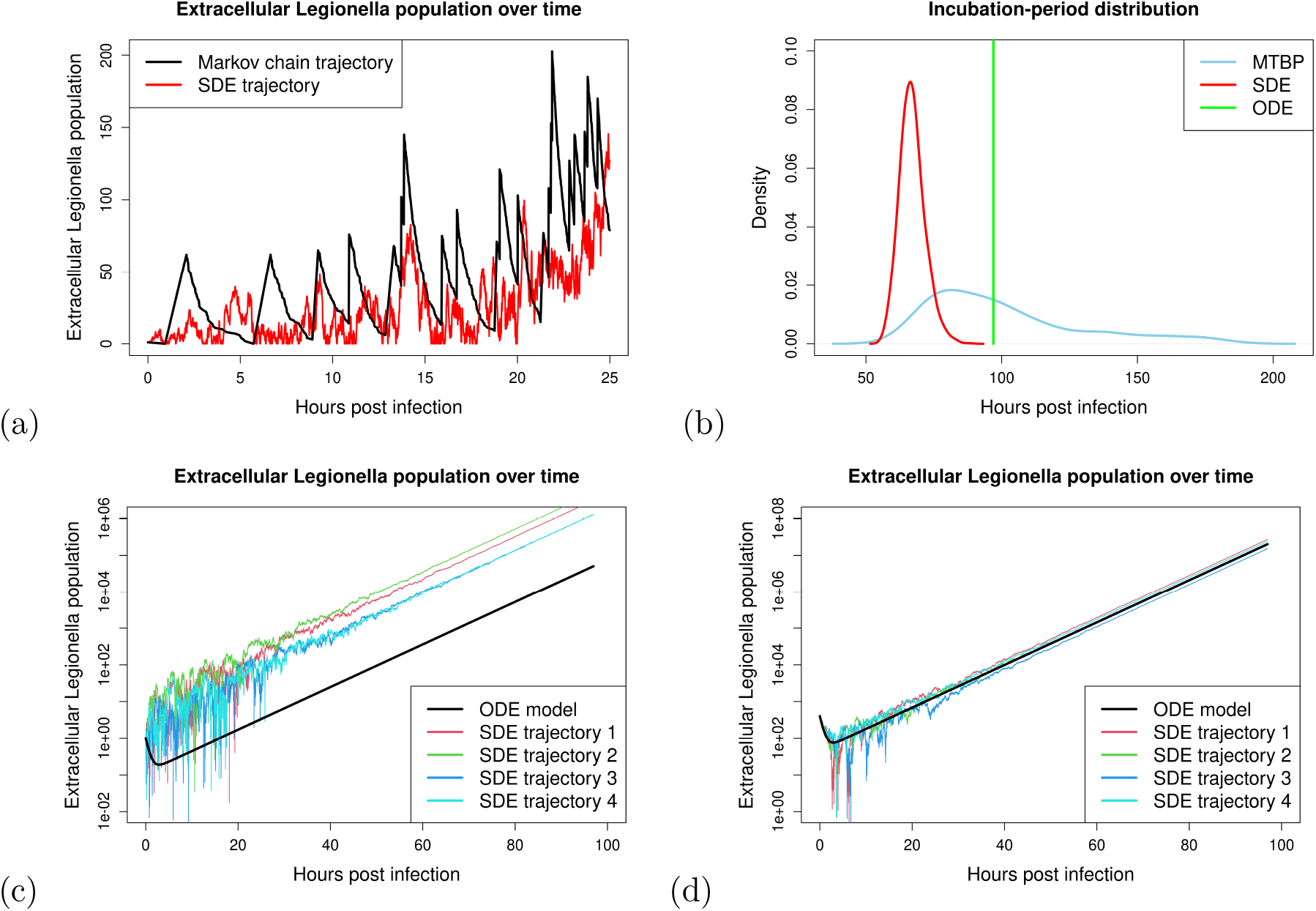
Plots of important features obtained when running all within-host models immediately after infection. Sub-figure 2(a) provides a visualisation of how SDE models fail to capture large jumps that occur in the extracellular Legionella populations following rupture events when the extracellular Legionella population is small. Sub-figure 2(b) provides incubation-period distributions from using the three within-host models for the full infection process. Sub-figure 2(c) provides a visualisation on the discrepancy between SDE and ODE models starting with one extracellular Legionella. Sub-figure 2(d) provides a visualisation of the discrepancy between SDE and ODE models beginning with a larger initial extracellular Legionella population (400 bacteria). Both sub-figures 2(c) and 2(d) are provided on the log scale.

In addition to this issue, different incubation-period distributions are obtained when estimating using various modelling techniques. For example, the deterministic ODE provides a point-mass incubation period that is larger than the mode obtained from both of the discrete-event stochastic and Gaussian SDE models. This discrepancy indicates potential issues with certain mathematical models in describing the infection process immediately after infection (Fig. 2b). To further compare the ODE and Gaussian SDE models, samples from the noise term **B**(*t*)*d****W*** _*t*_ may be either positive or negative when simulating the Gaussian SDE. In the case where the noise is negative, the trajectories are more likely to fade out. This fadeout typically leads to the process becoming extinct. Therefore, we observe a survivor bias. Specifically, surviving SDE trajectories are those that experience large jumps early on, which leads to their accelerated growth and shorter incubation periods (Fig. 2c). These jumps have a 1greater influence in the early stages when the extracellular *Legionella* population is smaller.

Therefore, simulating a Gaussian SDE with a larger initial condition (e.g., 400 extracellular *Legionella*) would result in reduced stochasticity and a model closer to the corresponding ODE than if a smaller initial condition was used (e.g., one extracellular *Legionella*) (Fig. 2c-d).

### 2.7. Model parameter estimates

Having derived the various within-host models, we now turn to the discussion of the parameter estimates to be used in these models. For the models outlined in Sections 2.1-2.5, parameter estimates are obtained from [16]. An experimental dataset containing the rupture times of bone-marrow derived macrophages from mice infected with *Legionella* [21] is used to estimate *λ* [15]. *λ* is defined as the reciprocal of the median rupture time of this dataset and is estimated as *λ* = 0.041 per hour. Moreover, an experimental dataset was considered in which human-derived macrophages were infected with *Legionella*. At 1, 24, 48 and 72 hours post-infection, the average number of intracellular *Legionella* within macrophages was recorded [24]. In [16], a logistic growth curve was fitted to this dataset, and a rupture size *G* was calculated as the value of the logistic growth curve at the median rupture time 1*/λ*. With a distribution for *λ* obtained through bootstrapping methods, a distribution of rupture sizes was obtained by iteratively sampling an estimate of *λ* and recalculating the rupture size through the logistic growth curve. This approach resulted in estimates of *G* = 62.792 *Legionella* and *r* = 32.072. Following this, an experimental dataset [12], in which the number of *Legionella* within the lungs of mice was recorded at intervals of 2, 24, 48 and 72 hours post-infection, was used to estimate *α* and *β*. The deterministic model described in Section 2.5 was fitted to this data, yielding estimates of *α* = 0.089 per hour and *β* = 1.088 per hour were obtained.

### 2.8. First passage times

In this section, we define two first passage times for our hybrid framework. We begin by defining a first passage time 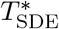, the time at which a Gaussian SDE begins to approximate the MTBP dynamics. Following this, we define another first passage time 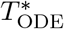, the time at which an ODE begins to approximate the MTBP dynamics.

To begin, we define the *first passage time* 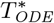 as the earliest time at which the Gaussian SDE model (developed in Section 2.4) provides an accurate approximation of the MTBP. More precisely, this transition occurs when the following two conditions are satisfied. The first condition is that the instantaneous probability of extinction *p*_SDE_(*t*), calculated from the Gaussian SDE model, must converge to the instantaneous probability of extinction *p*_BP_(*t*), calculated from the MBTP model, within a tolerance 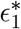. Mathematically, we require 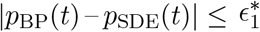 for 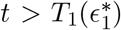. The SDE model’s extinction probability aligning with that of the MBTP model indicates that the SDE has successfully captured a fundamental property of the stochastic process. The second condition is that the coefficient of variation *c*_SDE_(*t*) of the Gaussian SDE model must converge to the coefficient of variation *c*_BP_(*t*) of the MBTP model, within a tolerance 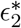. Mathematically, we require the first time max 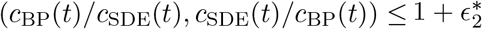 for 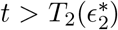. By requiring both models’ coefficients of variation to converge, we ensure that the SDE model is accurately capturing the level of variability or uncertainty present in the MBTP model. Having both of these conditions satisfied, we can guarantee a level of consistency between the MTBP and Gaussian SDE in terms of mean growth, extinction probability, and system fluctuation. We assume that consistency between these properties of the MTBP and Gaussian SDE models is sufficient for the Gaussian SDE model to approximate the MTBP dynamics.

Similarly, we define the *first passage time* 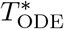 as the earliest time at which the ODE model (developed in Section 2.5) provides an accurate approximation of the MTBP. More precisely, this transition occurs when both of the following two conditions are satisfied. The first condition is that the probability of extinction *p*_BP_(*t*) of the MTBP at time *t* must be negligible within a tolerance 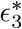. Mathematically, we require 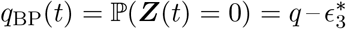 for 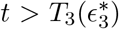. The instantaneous probability of extinction *p*_BP_(*t*) becoming negligible indicates that extinction within the MTBP model is an unrealistic outcome. This feature is consistent with the extinction probability of an ODE model. The second condition is that the extra-cellular *Legionella* and infected macrophage populations must grow according to the mean growth curve, within a tolerance 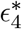. Mathematically, we require 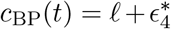 for 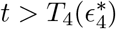.

By requiring the MTBP’s coefficient of variation to become constant, we find a point in time when the fluctuations are small enough to be negligible and system’s dynamics are governed primarily by the deterministic growth. We assume that consistency between these properties of the MTBP and ODE models is sufficient for the ODE model to approximate the MTBP dynamics.

From the two conditions required for 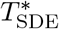, we define 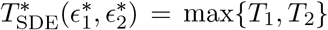. We assume that at 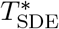, the expected size of each population 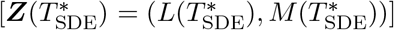 is an absorbing state for the MTBP. In this scenario, we may simulate the MTBP until absorption to generate a distributional estimate of 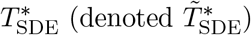. Similarly, from the two conditions required for 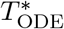, we define 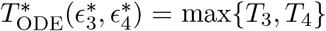. We assume that at 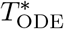, the expected size of each population 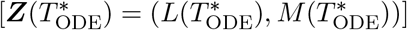 is an absorbing state for the MTBP (or Gaussian SDE starting from a sample of 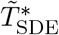 with initial condition 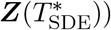. In this scenario, we may simulate the MTBP (or Gaussian SDE from a sample of 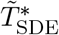 ) until absorption to generate a distributional estimate of 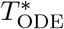 (denoted 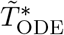).

Notably, all definitions (and consequently estimates) of *T*_*l*_ for *l* ∈ {1, 2, 3, 4} depend on the corresponding tolerance 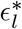 for *l* ∈ {1, 2, 3, 4}. Choosing appropriate values for 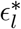 requires careful consideration, as a trade-off exists. For all *l* ∈ {1, 2, 3, 4}, *T*_*l*_ is a monotonically decreasing convex function of 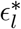. Therefore, setting 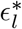 too small could cause a false conclusion that more time is required before the Gaussian SDE and ODE models can provide an accurate approximation of the MTBP dynamics. This false understanding would wrongly render the hybrid model unnecessary. In contrast, setting 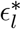 too large could cause the Gaussian SDE and ODE models to be applied prematurely, before they provide an accurate approximation of the MTBP dynamics. This premature application would introduce bias into our incubation-period estimates. The bias would appear as an incubation-period estimate larger that it would be if the ODE model was applied at an appropriate time. Therefore, choosing a different 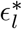 leads to different values of *T*_*l*_, and we must make a subjective decision about an acceptable tolerance *ϵ*_*l*_.

A common approach to find the *elbow point* in various trade-off problems is to find the optimal value of 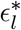 in which the curvature of 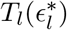 is maximized [19]. To balance the trade-off when determining 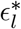 for *l* ∈ {1, 2, 3, 4}, we vary the value of *ϵ*_*l*_ in a reasonable range (we deem *ϵ*_*l*_ ∈ {10 ^− 5^, 10 ^− 1^} to be reasonable). With this range of *ϵ*_*l*_, we first calculate the corresponding *T*_*l*_ for each *ϵ*_*l*_. In this paper, we consider *T*_*l*_ as a function of *ϵ*_*l*_ and take our chosen value to be 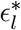 such that the gradient 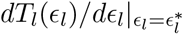 equals the average gradient over the domain, (*T*_*l*_(10 ^− 1^) – *T*_*l*_(10 ^− 5^))*/*(10 ^− 1^ – 10 ^− 5^). Estimating 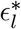 through this average-gradient approach corresponds to the point of maximum curvature [19].

## 3. Results

In this section, we provide the results from analysing the probability of extinction and the coefficient of variation of the various models developed in Section 2. First, we provide point estimates for 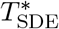 and 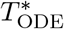. Second, we calculate the corresponding population thresholds, 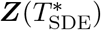 and 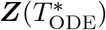, using the deterministic model. Third, we present the results from the within-host simulations of the MTBP and jump SDE models, which provide the distribution of 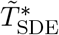. Fourth, we present the results from the within-host simulations of the MTBP, jump SDE, and Gaussian SDE models (starting from 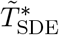 in the latter case), which provide the distribution of 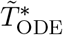. Fifth, we analyse the low-dose (initial condition of one extracellular *Legionella*) incubation period of Legionnaires’ disease, comparing the distributions obtained from the hybrid models with the low-dose incubation period derived from the MTBP model.

### 3.1. First passage times, 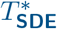 and 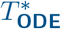

We calculate *T*_1_ and *T*_2_ based on the conditions defined in Section 2.8 (Fig. 3a-b). Using the elbow-finding procedure, we obtain 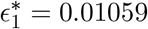 and *T*_1_ = 3.7 hours (Fig. 3a). Additionally, we find 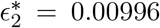 and *T*_2_ = 45.1 hours (Fig. 3b). Using the estimates for *T*_1_ and *T*_2_, we calculate 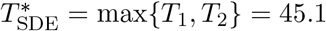 as a central estimate of the first passage time at which the Gaussian SDE model starts to approximate the MTBP model dynamics.

**Figure 3.**
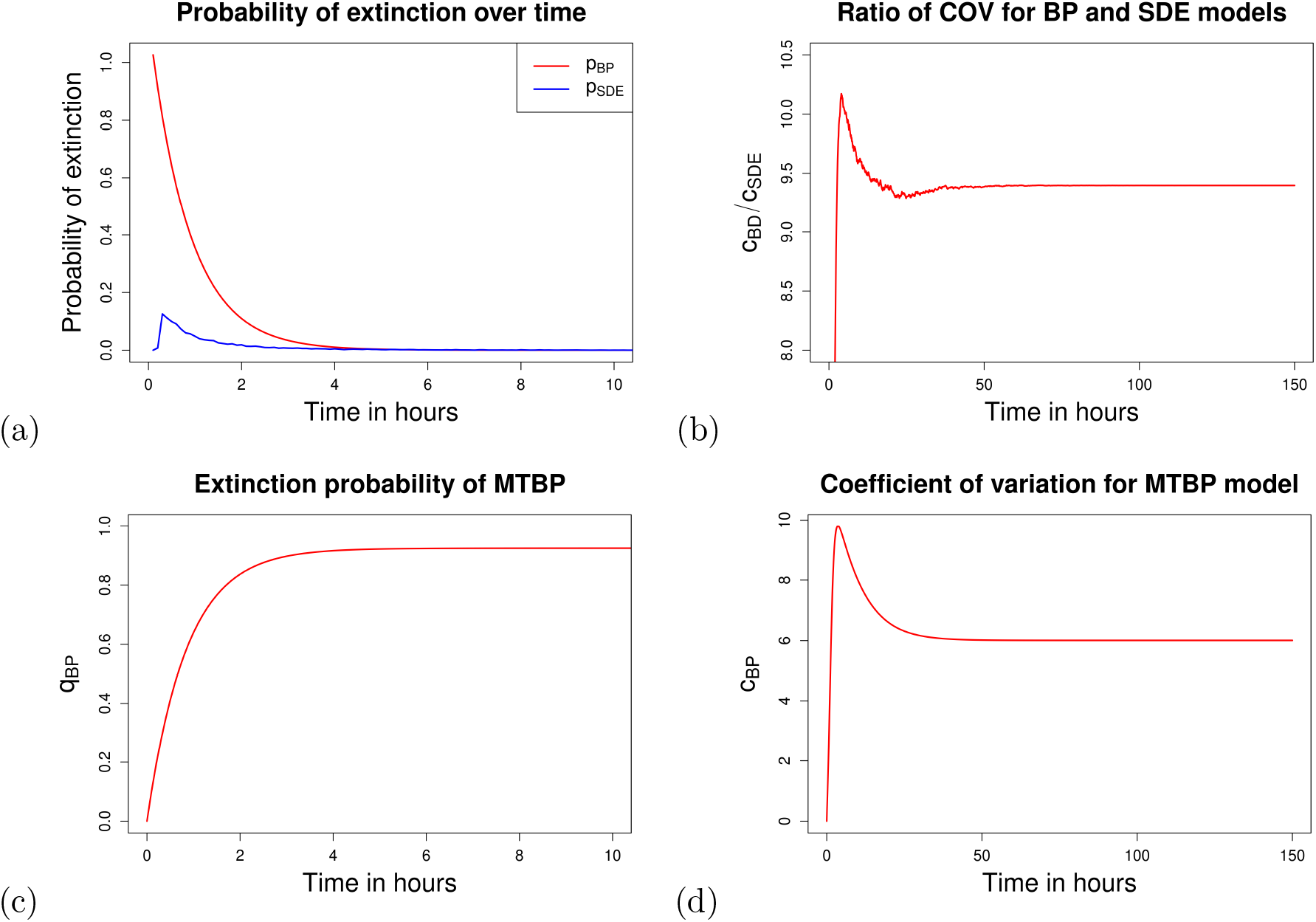
Plots for estimating 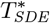 and 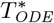. Sub-figure 3(a) provides a visualisation of how the probability of extinction changes over time for both the MTBP and Gaussian SDE models. Sub-figure 3(b) provides a visualisation of how the ratio of the coefficient of variation (COV) between the MTBP and Gaussian SDE models varies over time. Sub-figure 3(c) provides a visualisation of of the cumulative probability of extinction for the MTBP model increases over time. Sub-figure 3(d) provides a visualisation of how the coefficient of variation for the MTBP model varies over time.

Additionally, we calculate *T*_3_ and *T*_4_ based on the conditions defined in Section 2.8 (Fig. 3c-d). Using the elbow-finding procedure, we obtain 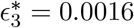 and *T*_3_ = 6.0 hours (Fig. 3c). Additionally, we find 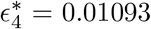 and *T*_4_ = 49.9 hours (Fig. 3d). Using these estimates for *T*_3_ and *T*_4_, we calculate 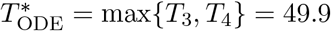 hours as the central estimate of the first passage time at which the ODE model starts to approximate the MTBP dynamics. These results indicate that the Gaussian SDE model becomes accurate only 4.8 hours earlier than the ODE model.

### 3.2. First passage time distributions, 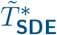 and 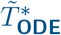

Following the analysis to estimate 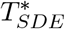, we calculate the absorbing populations, 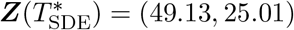 that are used within the MTBP and jump SDE models. We employ the Gillespie and Euler–Maruyama algorithms, respectively, to estimate 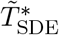 from 10,000 realizations of the simulations (Fig. 4). Similarly, we calculate the absorbing populations, 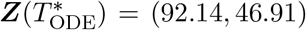 that are used within the MTBP, jump SDE and Gaussian SDE models. The Gillespie algorithm is used to simulate the MTBP model, whereas the Euler–Maruyama algorithm is used to simulate the SDE models. We then estimate 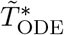 from 10,000 realizations of the simulations (Fig. 4).

**Figure 4.**
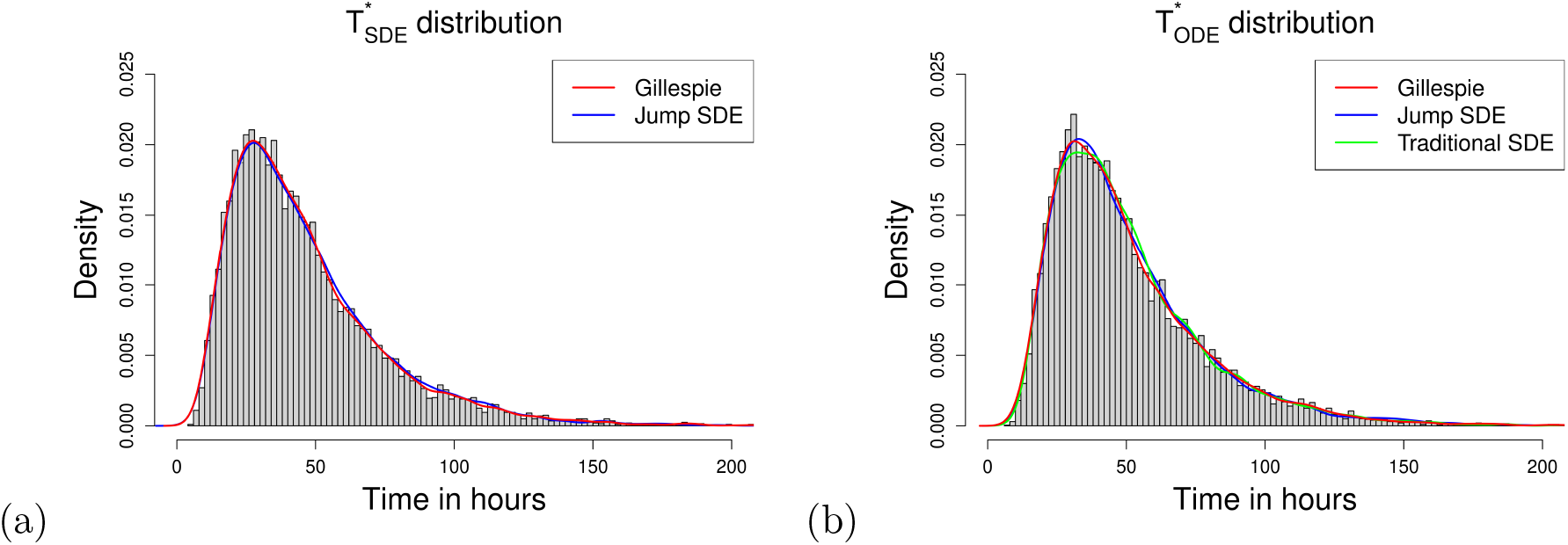
Plot of the distributions of 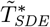 and 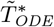. In red, we provide the kernel density obtained from running the MTBP model 10,000 times using the Gillespie algorithm. In blue, we provide the kernel density obtained from running the jump SDE 10,000 times using the Euler–Maruyama algorithm. In green, we provide the kernel density obtained from simulating the Gaussian SDE from 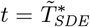 to 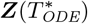 10,000 times using the Euler–Maruyama method.

Simulating the jump SDE until the process reaches 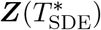 yields similar results to the MTBP for the distribution of 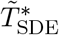 (Fig. 4a). These results support the validity of the jump SDE as a model that can approximate the MTBP dynamics from *t* = 0. The same conclusion can also be made from analysing the results of simulating the jump SDE from *t* = 0 until the process reaches 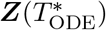 (Fig. 4b). Furthermore, simulating the Gaussian SDE from a sample of 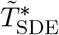 until the process reaches 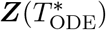 also yields similar results to the MTBP model for the distribution of 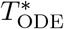 (Fig. 4). These results indicate that the conditions for 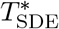 defined in Section 2.8 are reliable. Notably, the distributions for 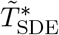 and 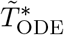 appear similar (Fig. 4a-b). Therefore, for our within-host model, the Gaussian SDE is accurate only for a short time before the ODE model becomes the appropriate modelling approach.

### 3.3. Low-dose incubation period of Legionnaires’ disease

From 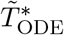, the extracellular *Legionella* and infected macrophage populations follow the exponential growth curve defined by (2.7). Therefore, from 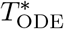 the within-host dynamics of Legionnaires’ disease may be modeled with a deterministic ODE. Solving the ODE model with initial condition 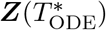 for the time 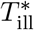 at which illness occurs, we obtain an estimate for 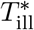 to be 96.9 hours. Further, we may calculate the time difference 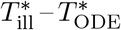. This difference is the duration during which the ODE is applied. We may use this time difference to obtain a distribution of the low-dose incubation period of Legionnaires’ disease (Fig. 5).

**Figure 5.**
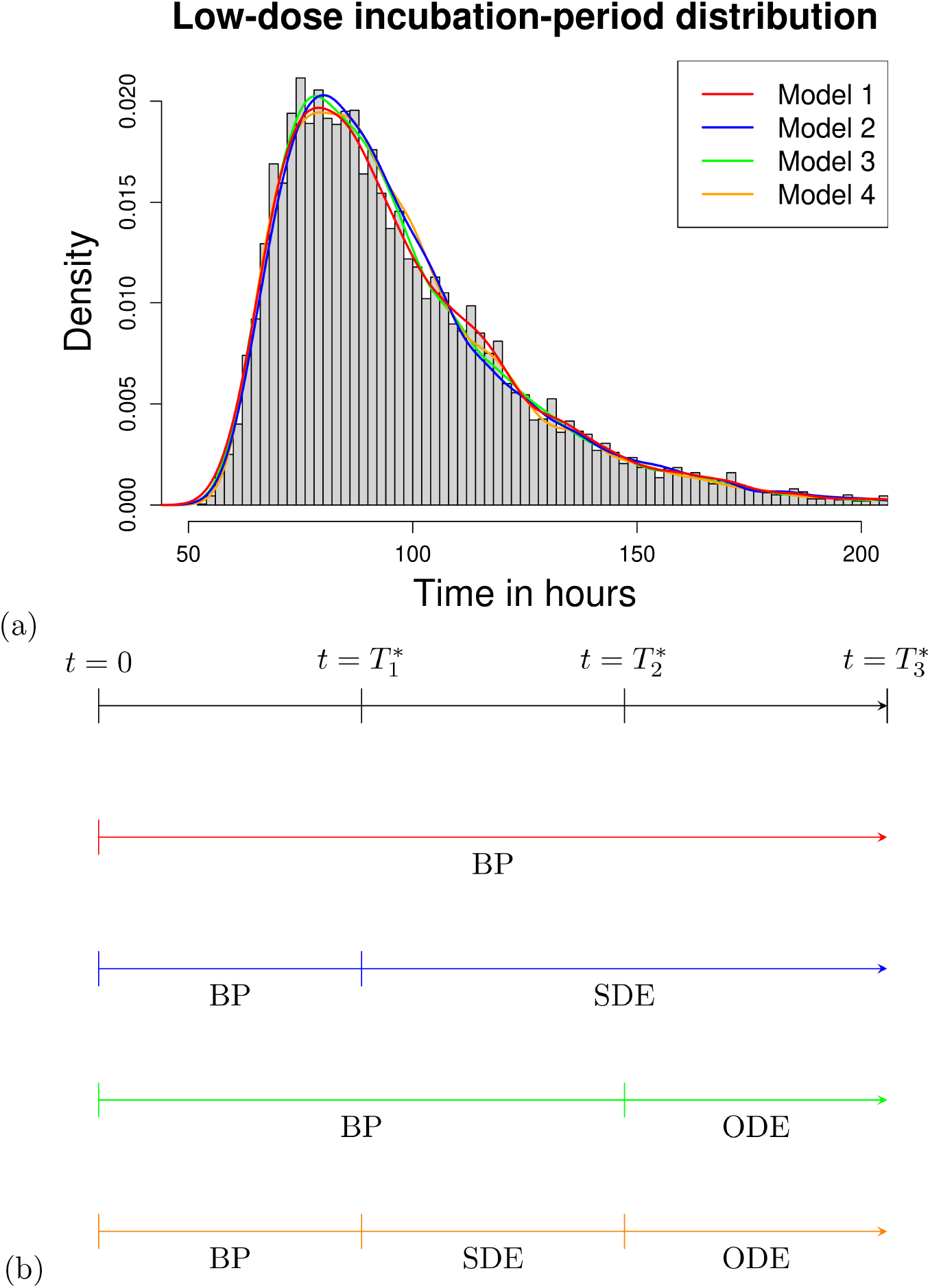
Plot of the estimated low-dose incubation period of Legionnaires’ disease. Model 1 represents the MTBP model simulated from t = 0 until 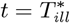. Model 2 represents the MTBP model simulated from t = 0 to 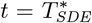, then the Gaussian SDE simulated from 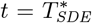 to 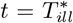. Model 3 represents the MTBP model simulated from t = 0 to 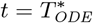, then the ODE model simulated from 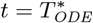 to 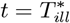. Model 4 represents the MTBP model simulated from t = 0 to 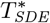, then the Gaussian SDE model simulated from 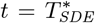 to 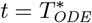, then the ODE model simulated from 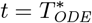 to 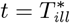.

We provide four approaches for obtaining a low-dose incubation-period distribution of Legionnaires’ disease using our first passage times 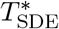 and 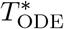 (Fig. 5). In the first instance, we simulate the MTBP model until a threshold *L*_*T*_ of extracellular *Legionella* are present within the lungs, following [16]. For the next approach, we simulate the MTBP model until the process reaches the absorbing state 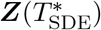. At this point, we run the Gaussian SDE model until *L*_*T*_ extracellular *Legionella* are present within the lungs. Moreover, in a third approach, we simulate the MTBP model until the process reaches absorbing state 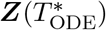.

At this point, we run the ODE model until *L*_*T*_ extracellular *Legionella* are present within the lungs. For our final approach, we simulate the MTBP model until the process reaches the absorbing state 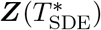. At this point, we run the Gaussian SDE model until the process reaches the absorbing state 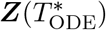. Then, we run the ODE model until *L*_*T*_ extracellular *Legionella* are present within the lungs. The third and fourth models represents our first and second hybrid frameworks presented in Fig. 5.

From Fig. 5, all four approaches provide very similar low-dose incubation-period distributions. Therefore, these simulations indicate that both first passage times 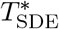 and 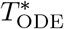 provide valid estimates for the time at which the Gaussian SDE and ODE models can be applied.

## 4. Discussion

In this paper, we introduced two novel hybrid approaches to model the within-host dynamics of Legionnaires’ disease. These models combine stochastic and deterministic methods, enabling a more accurate representation of the progression of infection. The MTBP-to-ODE hybrid framework is uncommon within mathematical modelling of bacterial dynamics. In addition to the implementation of this framework, this work introduces a further innovation: extending the hybrid model to incorporate an intermediary stage, modelled using a Gaussian SDE. This extension adds flexibility and accuracy to the model, allowing the model to capture a wider range of population dynamics. The ability to represent these intermediary dynamics is especially important in biological systems where transitions between stochastic and deterministic behaviour are gradual rather than abrupt.

To fully develop our hybrid models, we quantified the first passage times, 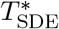 and 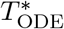, to describe when SDE and ODE models, respectively, become accurate approximations of the MTBP dynamics. We assumed that SDE and ODE models become appropriate approximations when two key properties converged to that of the MTBP model: first, the probability of extinction; and second, the coefficient of variation. Although our conditions for the ODE approximation have been applied in outbreak modelling [5], the conditions for switching to the Gaussian SDE model are new innovations developed in this research. Our results demonstrate that our choice of conditions for 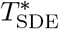 are reasonable, validating our inclusion of an intermediary stage in the hybrid model.

A critical aspect of this research was the data-driven method used to determine 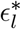. In contrast to previous studies where 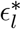 was chosen arbitrarily [5], our elbow-finding procedure ensures that the transition point is optimally chosen. An arbitrary choice could lead to the simpler model being before it provides an accurate approximation. Additionally, an arbitrary choice could result in the simpler model being applied later than it becomes approximate, falsely rendering the hybrid framework redundant. In this case, an incorrect conclusion could be made that the hybrid approach is redundant. Our approach for determining the ideal is 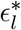 [19] is mathematically valid and provides us with a method to estimate *T*_*l*_ without the two risks described above. Noticing the similar low-dose incubation-period distributions estimated from different modelling approaches (Fig. 5) provides credibility for method in setting 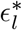.

Using our fitted hybrid models, we generated estimates of the incubation period for Legionnaires’ disease, which ranged from two to ten days for low-dose infections. This result aligns closely with reported data from human outbreaks in the literature [4, 6, 8, 10, 11, 13, 15, 16]. This consistency with empirical data reinforces the validity of our hybrid models, suggesting that they provide a reliable and flexible framework for accurately describing the within-host dynamics of Legionnaires’ disease during different phases of the infection.

In summary, we developed two hybrid models that provide a more precise and computationally efficient way to simulate the within-host dynamics of Legionnaires’ disease. By leveraging both stochastic and deterministic models, our models have provided valuable insights into the disease’s progression and incubation period. Importantly, these models address key limitations in previous work [16] by incorporating intermediary dynamics. As a result, our hybrid models offer a valuable tool to quickly and accurately model the within-host dynamics of Legionnaires’ disease. This increased speed may allow one to test more complex modeling assumptions, which would otherwise be computationally infeasible.

Future research could explore extending these hybrid models to account for additional complexities, such as the adaptive immune response or variations in pathogen virulence. Additionally, further work may apply this framework to other pathogens to demonstrate its generalisability across different diseases. Ultimately, our hybrid framework represents a step forward in modelling within-host dynamics, providing a foundation for more accurate and efficient simulations. This may allow researchers to test model assumptions and obtain key results such as the dose–response or incubation period of diseases in a more computationally feasible manner.

## Acknowledgments

NJ acknowledges support from the Engineering and Physical Sciences Research Council (EPSRC) and Mathematics and Data in Scientific and Industrial Modelling (MADSIM) at the University of Manchester for funding of their studentship.

IH was supported by the JUNIPER modelling consortium (grant MR/V038613/1) the National Core Study on Transmission (PROTECT) and by the UKRI Impact Acceleration Account (IAA 386). NJ and IH also acknowledge the UK Health Security Agency (UKHSA) for honorary contracts and funding (for IH). The views expressed are those of the author(s) and not necessarily those of the Department of Health or UKHSA.

The authors have made the code and data available at: https://github.com/NyallJamieson/Hybrid-paper

## Notes

**Funding:** NJ acknowledges support from the Engineering and Physical Sciences Research Council (EPSRC) and Mathematics and Data in Scientific and Industrial Modelling (MADSIM) at the University of Manchester for funding of their studentship.

### Competing Interest Statement

The authors have declared no competing interest.

